# *In Vivo* Surface Reconstruction of Three Signal-defined Intracortical Layers Using 5T 3D FLAIR MR Imaging

**DOI:** 10.64898/2026.02.05.703937

**Authors:** Shui Cao, Lu Shi, Jiameng Liu, Zifeng Lian, Lin Teng, Qin Xiao, Xiaona Xia, Kaicong Sun, Feng Shi, Xiangshui Meng, Dinggang Shen

**Affiliations:** School of Biomedical Engineering & State Key Laboratory of Advanced Medical Materials and Devices, ShanghaiTech University, Shanghai, China; Department of Radiology, Qilu Hospital, Cheeloo College of Medicine, Shandong University, Jinan, China; Department of Radiology, Qilu Hospital (Qingdao), Cheeloo College of Medicine, Shandong University, Qingdao, China; United Imaging Intelligence, Shanghai, China; Shanghai Clinical Research and Trial Center, Shanghai, China

**Author notes:** These authors contributed equally to this work and are co-first authors (Shui Cao, Lu Shi).

## Abstract

**Background:** *In vivo* whole-cortex quantification of intracortical signal-defined layering on the routinely acquired structural MRI remains limited.

**Purpose:** To develop and validate an automated framework to reconstruct three intracortical signal-defined layers from 5T three-dimensional (3D) T2-weighted fluid-attenuated inversion recovery (FLAIR) and to characterize whole-cortex morphometrics and regional organization across a prespecified cortical organizational framework.

**Materials and Methods:** In this retrospective study, 5T 3D FLAIR images were acquired between February and July 2024. Brain Multi-Layer Surface Reconstruction (BrainMLSR) reconstructed three intracortical signal-defined layers, and derived intracortical layer thickness and surface area measures and ratios. Performance was evaluated against manual annotations and assessed for test-retest repeatability (n=13) and cross-site feasibility (n=2). Paired two-tailed *t*-tests and linear mixed-effects models were used. A proof-of-concept analysis compared Heschl’s gyrus ratios between 19 patients with temporal lobe epilepsy (TLE) and 19 age-matched healthy controls (HC).

**Results:** A total of 270 healthy participants (mean age, 54.4±14.5 years; 146 men) were included. Agreement with manual hypointense-layer annotations was high (Dice, 0.960±0.003), and was similar in the cross-site dataset (Dice, 0.954±0.009). In the test-retest dataset, average symmetric surface distance was less than 0.1 mm. Across prespecified systems, thickness and surface area ratios varied by region; within an auditory-perisylvian hierarchy, banksSTS showed a localized turning point with an increased hyperintense layer thickness ratio and decreased hypointense layer thickness ratio, accompanied by inflections in surface area ratios (*P* < .001). In bilateral Heschl’s gyrus, hypointense (left: 0.619±0.262 *vs* 0.881±0.102; right: 0.607±0.310 *vs* 0.907±0.141 mm) and isointense (left: 0.406±0.225 *vs* 0.678±0.128; right: 0.478±0.232 *vs* 0.808±0.176 mm) layer thicknesses were lower in TLE than in HC (all *P*<.001).

**Conclusion:** BrainMLSR enabled accurate and repeatable *in vivo* reconstruction of three intracortical signal-defined layers from a single 5T 3D T2-weighted FLAIR acquisition and provided whole-cortex boundary-based morphometry with interpretable regional organization.

**Key Results:**

1. In this retrospective study of 270 participants, Brain Multi-Layer Surface Reconstruction (BrainMLSR) showed high agreement with hypointense-layer annotations (Dice, 0.960±0.003).
2. Within an auditory-perisylvian hierarchy, the hyperintense-layer thickness ratio peaked (0.620±0.003) and the hypointense-layer ratio was the lowest (0.156±0.002) in the bank of the superior temporal sulcus (*P* < .001).
3. In a proof-of-concept analysis, patients with temporal lobe epilepsy (*vs* healthy controls) had higher hyperintense-layer thicknesses ratio in the left hemisphere (0.523±0.085 *vs* 0.417±0.036, *P*<.001).

**Summary:** An automated multiple-layer surface reconstruction framework (BrainMLSR), applied to 5T 3D T2-weighted FLAIR images, produced reproducible whole-cortex, signal-defined laminar morphometry and demonstrated coherent patterns across prespecified cortical organizational framework.

## INTRODUCTION

Cortical abnormalities do not always manifest as uniform global atrophy. Instead, many neurologic disorders are more likely to produce layer-selective injury or intracortical redistribution rather than homogeneous cortical thinning (1). Representative examples include subpial cortical demyelination in multiple sclerosis (2), dyslamination in focal cortical dysplasia (3), intracortical abnormalities in focal epilepsy (4,5), and layer-preferential pathologic distributions across neurodegenerative disorders (6). Developing layer-specific, repeatable quantitative measures may therefore complement conventional morphometric summaries such as mean cortical thickness or volume.

MRI approaches have been developed to study intracortical layer organization *in vivo* (7). Across cortical depth, MRI intensity profiles have been linked to the underlying myeloarchitecture and cytoarchitecture (8). Using 7T T2-weighted fluid-attenuated inversion recovery (FLAIR), prior work reported intracortical banding with a three-band appearance in the examined cortical regions (9). At 5T, three-dimensional (3D) T2-weighted FLAIR provides intracortical contrast that can be separated into three signal-defined layers in signal profiles: a superficial hyperintense layer adjacent to the pial surface, an intermediate hypointense layer, and a deep isointense layer bordering the white matter surface. These signal-defined layers provide a substrate for noninvasive intracortical layer morphometrics.

However, quantification of signal-defined intracortical layer measures remains limited. Many MRI approaches summarize intracortical organization as line-based or region-based depth profiles, restricting spatial coverage and complicating surface-based, whole-cortex testing (8-10). Moreover, common surface pipelines sample signals using geometry-derived intracortical coordinates (e.g., equidistant or equivolume surfaces); while useful for depth-based sampling, these interpolated surfaces do not represent signal-defined boundary surfaces (11-15). Although a clinically familiar structural sequence such as FLAIR can encode intracortical contrast in its signal profiles, a standardized framework to convert this contrast into participant-level, whole-cortex intracortical layer morphometrics remains lacking.

To address this gap, we developed Brain MultiLayer Surface Reconstruction (BrainMLSR), an automated framework that reconstructs three signal-defined intracortical layers from 5T 3D T2-weighted FLAIR and derives layer-specific morphometrics including thickness and surface area. A cortical organizational framework was prespecified, comprising key primary regions and two region-of-interest hierarchies designed to approximate a unimodal-to-association (transmodal) progression (16-19). The purpose of this study was to validate BrainMLSR measurement properties (manual-annotation agreement, test-retest reliability, and cross-site feasibility), characterize system-level spatial patterning within the prespecified framework, and, in an exploratory proof-of-concept analysis, assess whether BrainMLSR-derived metrics could capture subtle intracortical abnormalities in temporal lobe epilepsy without additional MRI-visible lesions beyond hippocampal sclerosis.

## MATERIALS AND METHODS

### Ethics Statement

This retrospective, multicenter study was approved by the institutional review boards of all participating centers, and informed consent was waived owing to the retrospective design.

### Participants

This study analyzed four 5T MRI datasets from two centers. The primary cohort comprised 270 neurologically healthy participants (mean age, 54.4±14.5 years; 146 men) recruited at Qilu Hospital (Qingdao) of Shandong University (Center 1) between February and July 2024. Additional datasets included: two participants from Shandong Provincial Third Hospital (Center 2) for cross-site feasibility; 13 participants from Center 1 with test-retest scans separated by an interval of approximately 15 minutes for reliability; and 19 patients from Center 1 with temporal lobe epilepsy (TLE) for exploratory assessment of clinical potential.

Healthy and TLE patients met their respective eligibility criteria; no subject overlap exists with any published, in-press, or under-review manuscripts. (Figure 1, Appendix S1).

**Figure 1.**
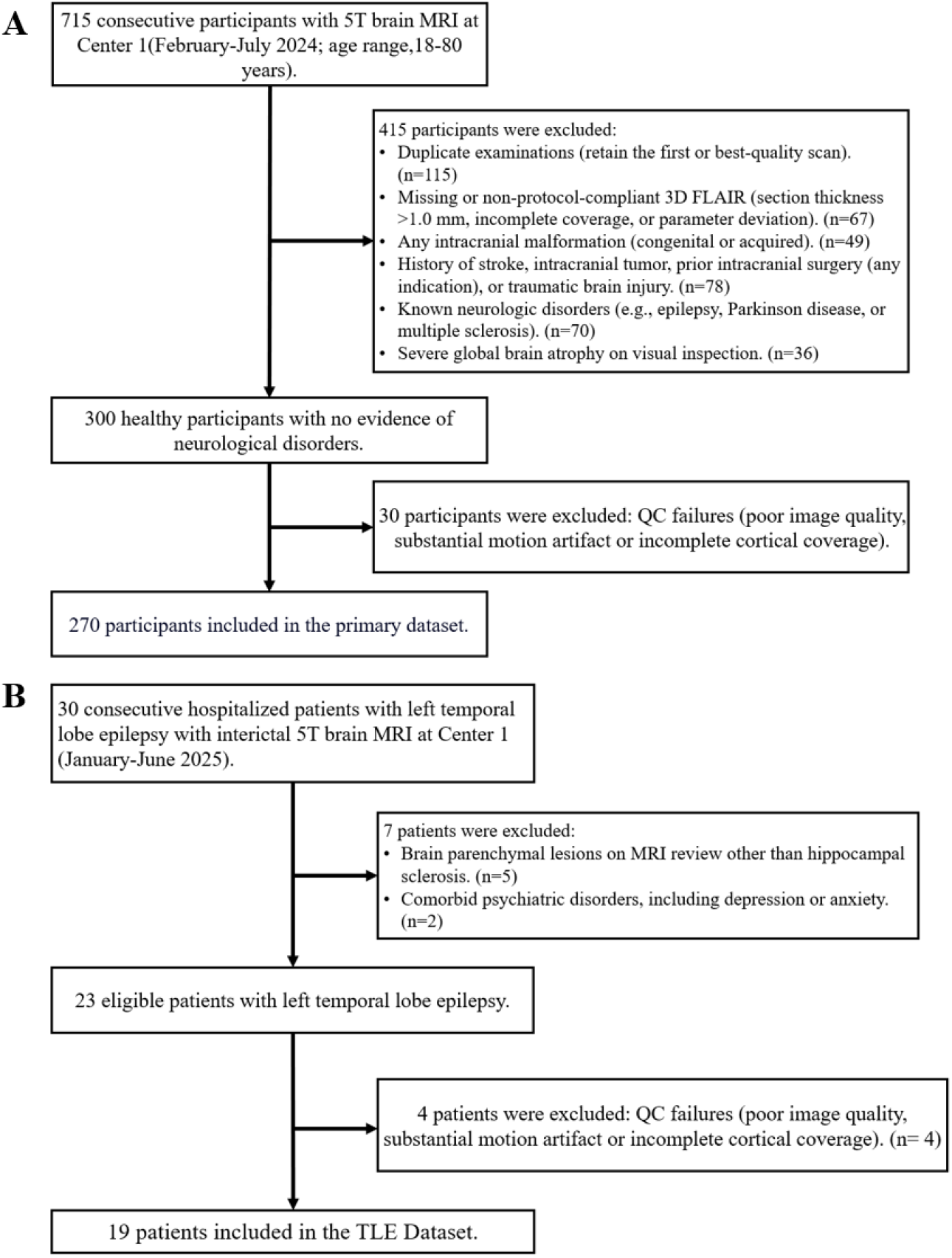
Participant selection flowchart illustrating inclusion and exclusion criteria. All excluded participants or patients had no usable data for any variable of interest. (A) The primary dataset (n = 270) is used for cortical surface reconstruction. Both the test-retest (n = 13) and cross-site (n = 2) datasets are processed through the same eligibility screening pipeline and data quality control (QC). TBI = Traumatic Brain Injury. (B) The temporal lobe epilepsy (TLE) dataset (n = 19) is used to demonstrate the potential clinical utility of our method in a proof-of-concept analysis.

### Imaging Protocol

At Center 1, 5T 3D T2-weighted FLAIR images were acquired (uMR Jupiter, United Imaging Healthcare; TR/TE/TI=6500/370/1950 ms; isotropic 0.533 mm^3^) across all datasets. For external validation, 5T 3D FLAIR images were acquired at Center 2 using comparable parameters (voxel size, 0.4×0.4×0.8 mm^3^). Acquisition details are provided in Appendix S2.

All images underwent visual inspection to exclude substantial motion or ghosting artifacts before surface reconstruction. Manual annotations of the hypointense layer and quality assessments of reconstructed surfaces were performed independently by two co-first authors (S.C. and L.S.)

### BrainMLSR Framework

The proposed BrainMLSR framework for multi-layer surface reconstruction is illustrated in Figure 2. The pipeline consists of three main steps.

**Figure 2.**
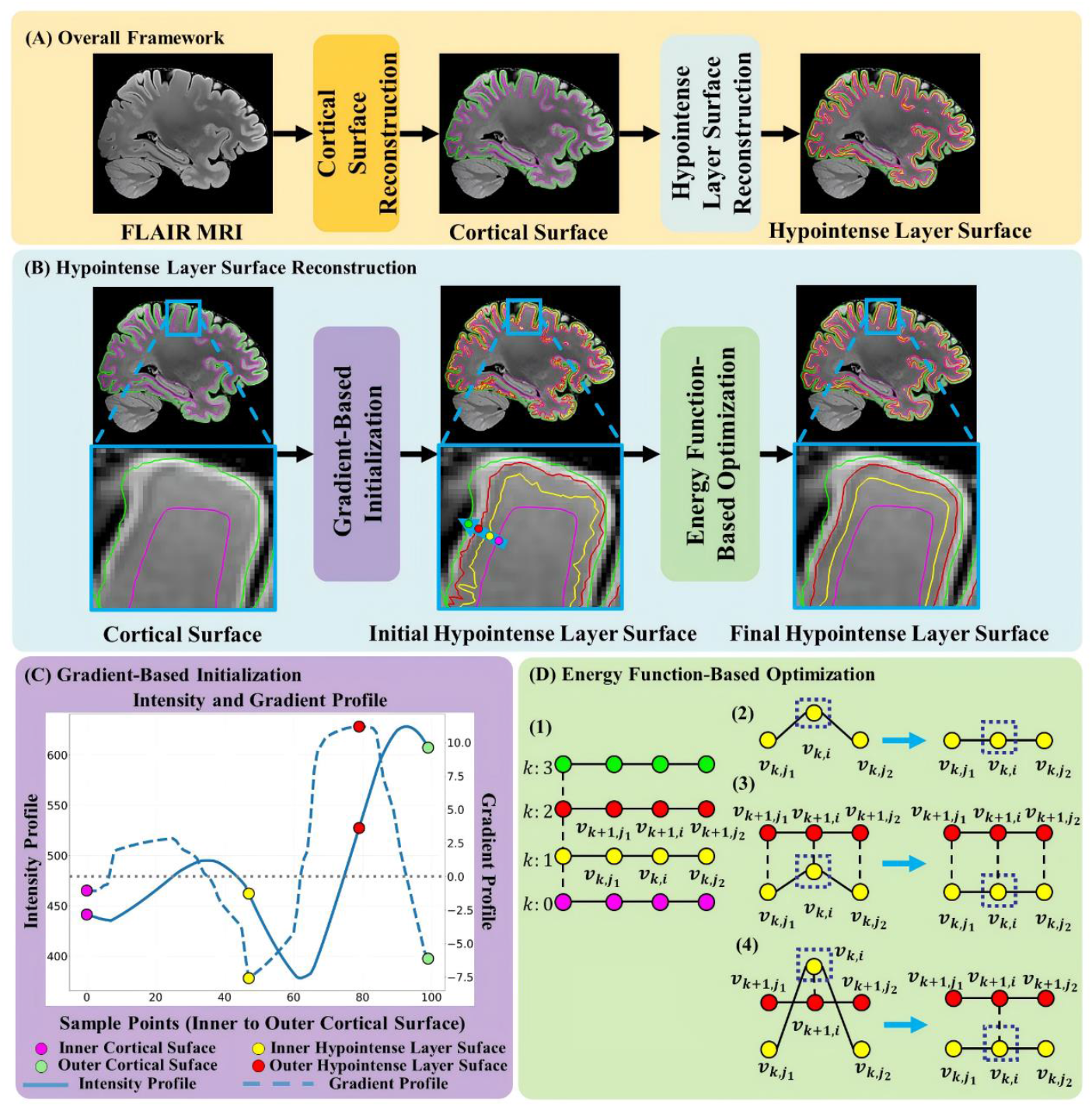
Overview of our BrainMLSR pipeline for hypointense layer surface reconstruction from 5T 3D T2-weighted FLAIR MR images (3D FLAIR). Images are from a 45-year-old man in the primary dataset. (A) 3D FLAIR MR images are preprocessed to reconstruct the pial and white matter surfaces, followed by hypointense layer surface reconstruction; (B) Initial hypointense layer surfaces are generated by gradient-based initialization and then refined via energy function-based optimization; (C) Signal intensity and gradient profiles sampled along the blue arrow in (B). (D) The components of the energy function-based optimization include: (D1) vertex models of the surfaces, (D2) surface smoothness constraint, (D3) surface distance consistency penalty, and (D4) surface intersection constraint. Abbreviations: BrainMLSR = Brain Multi-Layer Surface Reconstruction; FLAIR = fluid-attenuated inversion recovery; 3D = three-dimensional.

Step 1: Cortical surface reconstruction from MR images. First, 5T 3D FLAIR images are processed using our previous validated pipeline (20, 21) to obtain the inner and outer cortical surfaces (Figure 2A).

Step 2: Hypointense layer surface initialization. Next, the inner and outer surfaces of the hypointense layer are initialized based on intracortical intensity profiles (Figure 2B). Intensity profiles are sampled along surface normal from the inner to the outer cortical surfaces. As shown in Figure 2C, these profiles typically exhibit a downward trend toward the hypointense layer followed by an upward trend toward the hyperintense layer. From the corresponding gradient profiles, the point with the maximum positive gradient marks the outer surface of the hypointense layer, whereas the point with minimum negative gradient marks its inner surface. (Appendix S3)

Step 3: Energy function-based multi-surface optimization. Finally, the initially coarse hypointense layer surfaces are jointly refined using an energy function-based multi-surface optimization framework (Figure 2D). The surfaces of hypointense layer are iteratively refined while enforcing four key constraints to ensure biologically plausible and topologically correct reconstructions. 1) Surface smoothness promotes local geometric regularity and prevents spurious irregularities; 2) Inter-surface distance consistency maintains approximately uniform spacing between adjacent surfaces; 3) Image-gradient fidelity attracts vertices toward regions of strong intensity gradients corresponding to true layer boundaries; 4) Surface-intersection prevention ensures adjacent surfaces not to intersect or overlap. (Appendix S4)

### Technical Performance Evaluation

Surface reconstruction accuracy was evaluated in four participants from Center 1. For each participant, we generated two types of masks for the hypointense layer in selected local cortical patches: 1) image-based masks, manually tracing the hypointense band directly on FLAIR images, and 2) surface-guided masks, constrained to the region between the reconstructed inner and outer hypointense layer surfaces (Figure 3B). Spatial agreement between the two masks was quantified using the Dice similarity coefficient (DSC).

**Figure 3.**
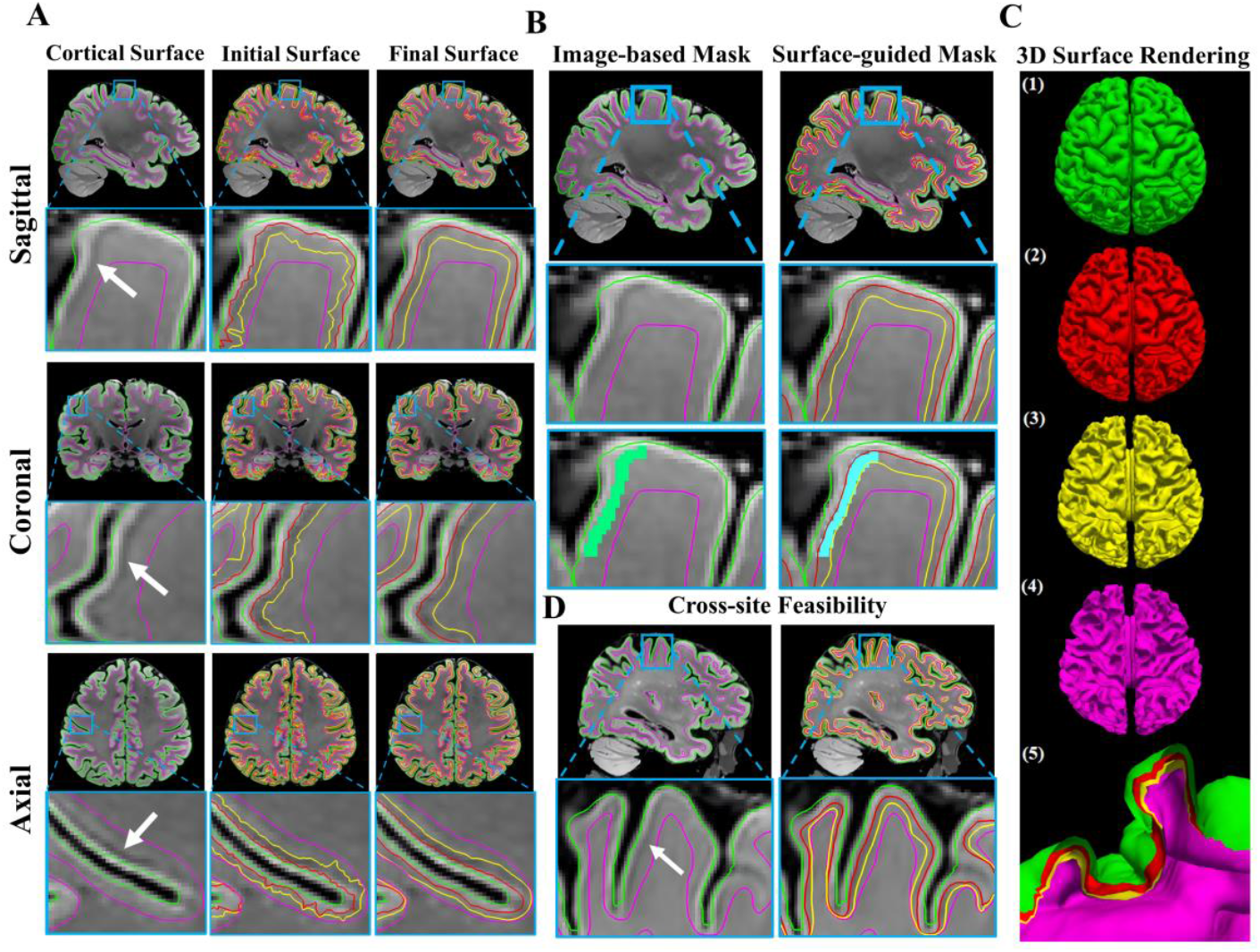
Qualitative evaluation and visualization of BrainMLSR results across imaging planes, hemispheres, and sites. (A) Reconstruction of multi-signal-layer cortical surfaces overlaid on 5T FLAIR MR images in sagittal, coronal, and axial views (a 45-year-old man from the primary dataset). The green, red, yellow, and purple surfaces represent the pial surface, the outer and inner surfaces of the hypointense layer, and the white-matter surface, respectively. White arrows indicate the hypointense layers. Representative regions are enlarged within blue boxes. (B) Illustration of manual masks used for accuracy evaluation (from the same participant as in (A)). The green mask denotes the image-based mask of the hypointense layer, and the blue mask represents the surface-guided mask derived from the surfaces of the hypointense layer. (C) 3D renderings of reconstructed multi-layer surfaces for both hemispheres: (C1) pial surfaces, (C2) outer surfaces of the hypointense layer, (C3) inner surfaces of the hypointense layer, (C4) white-matter surfaces, and (C5) all surfaces shown together to illustrate their nested configuration. (D) Demonstration of cross-site feasibility in a representative region from a 68-year-old man scanned at Center 2 using a 5T MRI scanner. Abbreviations: BrainMLSR = Brain Multi-Layer Surface Reconstruction; FLAIR = fluid-attenuated inversion recovery.

Cross-site feasibility was also evaluated by comparing image-based and surface-guided annotations in two participants from Center 2 using the DSC.

Test-retest reliability was assessed by applying BrainMLSR to the repeated scans for quantifying geometric consistency of hypointense-layer surfaces using both average symmetric surface distance (ASSD) and Hausdorff distance (HD).

### Statistical Analysis

Whole-hemisphere mean thickness and integrated surface area were computed for each intracortical layer per hemisphere. Inter-hemispheric differences and within-hemisphere inter-layer differences were assessed using two-tailed paired *t*-tests.

For regional analyses, we computed six layer-specific metrics: HyperR, HypoR, and IsoR (the thickness of the hyperintense, hypointense, and isointense layers, respectively, normalized by total cortical thickness) and PialR, OuterR, and InnerR (the surface areas of the pial, outer hypointense, and inner hypointense surfaces, respectively, normalized by white-matter surface area). These ratios were analyzed within each prespecified component of the cortical organizational framework using linear mixed-effects models, with cortical region, hemisphere, and age as fixed effects and participants a random intercept. To explore clinical utility, we compared layer-specific metrics in Heschl’s gyrus between TLE patients and healthy controls (HC) using a two-tailed Mann-Whitney U test.

All analyses were conducted in Python (version 3.8) by S.C. The sample size (n = 270) exceeds those of prior high-field MRI studies of intracortical architecture and provides adequate power for within-subject comparison. Statistical significance was defined as *P* < .05. (Appendix S6)

## Data Availability

The BrainMLSR code is publicly available at https://github.com/shuicao-water/BrainMLSR.git.

## RESULTS

### 1. Participant Characteristics

Of 300 neurologically healthy participants initially screened at Center 1, 30 were excluded due to poor image quality, as detailed in the study flow diagram (Figure 1A). The primary dataset comprised 270 healthy participants (146 men, 124 women; mean age, 54.4±14.5 years]; age range, 18-78 years) (Table 1).

**Table 1:** Participant Demographics of the Primary Dataset.

| Characteristic | All Participants (n = 270) | Men (n = 146) | Women (n = 124) |
| --- | --- | --- | --- |
| Sex, n (%) | - | - | - |
| Men | 146 (54.1%) | - | - |
| Women | 124 (45.9%) | - | - |
| Age, years | 54.4±14.5 | 53.5±14.7 | 55.4±14.2 |
| Age range, years | 18-78 | 18-78 | 19-77 |

### 2. Technical Performance

The BrainMLSR framework reconstructed cortical surfaces corresponding to three signal-defined intracortical layers (hyperintense, hypointense, and isointense layers) from 5T 3D FLAIR images and generated four boundary surfaces (pial surface, outer and inner surfaces of hypointense layer, and white-matter surface) (Figure 3A). 3D renderings of these surfaces, shown individually and in combination, illustrate their nested spatial configuration (Figure 3C).

Ablation experiments compared the initial surfaces generated by gradient-based initialization alone with the final surfaces produced after energy function-based optimization. The initial surfaces exhibited jagged or spiky profiles, whereas the optimized surfaces appeared smoother and more regular (Figure 3A). Additional qualitative examples from four subjects further illustrate the fidelity of the reconstructed hypointense layer surfaces, as shown in Figure 4.

**Figure 4.**
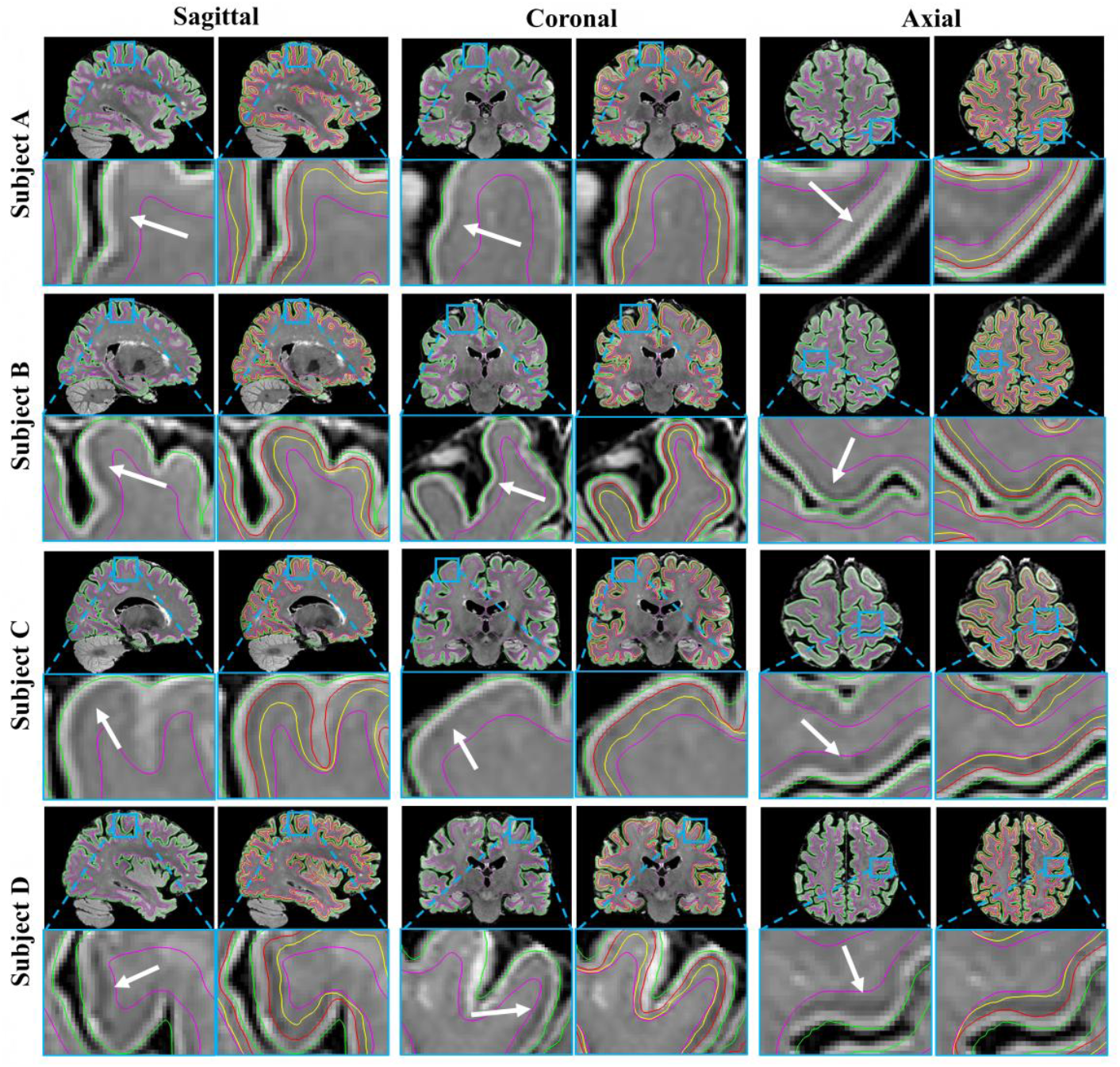
Reconstruction of three signal-defined intracortical layers in four neurologically healthy subjects from the primary dataset overlaid on 5T FLAIR MR images in sagittal, coronal, and axial planes (Subject A, 32-year-old woman; Subject B, 53-year-old woman; Subject C, 66-year-old man; Subject D, 77-year-old woman). The green, red, yellow, and purple surfaces represent the pial surface, outer and inner surfaces of the hypointense layer, and the white-matter surface, respectively. Arrows indicate the hypointense layers. Insets (blue boxes) show magnified views.

In four participants from the primary dataset, the DSC between image-based and surface-guided masks of the hypointense layer was 0.960±0.003. In two participants from the cross-site dataset, feasibility was assessed using the same annotation protocol, yielding a DSC of 0.954±0.009. (Figure 3D).

Reliability of hypointense layer surface reconstruction was assessed in 13 participants from test-retest dataset. ASSD values were submillimeter (<0.1 mm), and HD remained below 1.4 mm across all surfaces and hemispheres (Table 2).

**Table 2:** Test-Retest Reliability of BrainMLSR for Hypointense Layer (HL) Surfaces (ASSD and HD in mm)

| Surface | ASSD (mm, Mean ± SD) | HD (mm, Mean ± SD) |
| --- | --- | --- |
| Inner HL Surface (LH) | $0.093 \pm 0.009$ | $1.118 \pm 0.205$ |
| Outer HL Surface (LH) | $0.089 \pm 0.010$ | $1.068 \pm 0.188$ |
| Inner HL Surface (RH) | $0.093 \pm 0.008$ | $1.110 \pm 0.224$ |
| Outer HL Surface (RH) | $0.089 \pm 0.008$ | $1.334 \pm 0.114$ |

Note. The test-retest dataset and the same BrainMLSR pipeline are used to quantify differences between the inner and outer surfaces of the hypointense layer reconstructed from the two scan sessions, separately for the left (LH) and right (RH) hemispheres. Values are presented as mean ± standard deviation (SD) for both average symmetric surface distance (ASSD) and Hausdorff distance (HD), in millimeters (mm).

### 3. Whole-cortex Layer-specific Metrics

In the primary dataset, mean layer thickness was computed for the three signal-defined layers, and surface areas were computed for the four reconstructed boundary surfaces (Figure 5A, Table 3, Appendix Table 1).

**Table 3:** Layer-Specific Intracortical Cortical Thickness and Surface Area in Left and Right Hemispheres, with Interhemispheric Comparisons.

| Layer | LH (mm, Mean ± SD) | RH (mm, Mean ± SD) | *P* Value |
| --- | --- | --- | --- |
| Hyperintense Layer | 1.554 ± 0.106 | 1.551 ± 0.111 | .02 |
| Hypointense Layer | 0.626 ± 0.101 | 0.614 ± 0.101 | <.001 |
| Isointense Layer | 0.676 ± 0.074 | 0.664 ± 0.073 | <.001 |
| Total Cortex | 2.857 ± 0.097 | 2.828 ± 0.096 | <.001 |
| Surface | LH (cm<sup>2</sup>) | RH (cm<sup>2</sup>) | *P* Value |
| Pial Surface | 969.15 ± 82.07 | 975.75 ± 83.24 | <.001 |
| Outer HL Surface | 935.24 ± 84.83 | 942.66 ± 86.15 | <.001 |
| Inner HL Surface | 797.30 ± 78.45 | 802.80 ± 78.94 | <.001 |
| White Surface | 700.05 ± 70.73 | 705.06 ± 71.02 | <.001 |

Note. Values represent whole-hemisphere means ± standard deviation (SD) across all participants ( $n = 270$ ). *P* values correspond to paired two-tailed *t*-tests comparing left and right hemispheres without correction for multiple comparisons (significance threshold: $P < .05$ ). Abbreviations: LH = left hemisphere; RH = right hemisphere.

**Figure 5.**
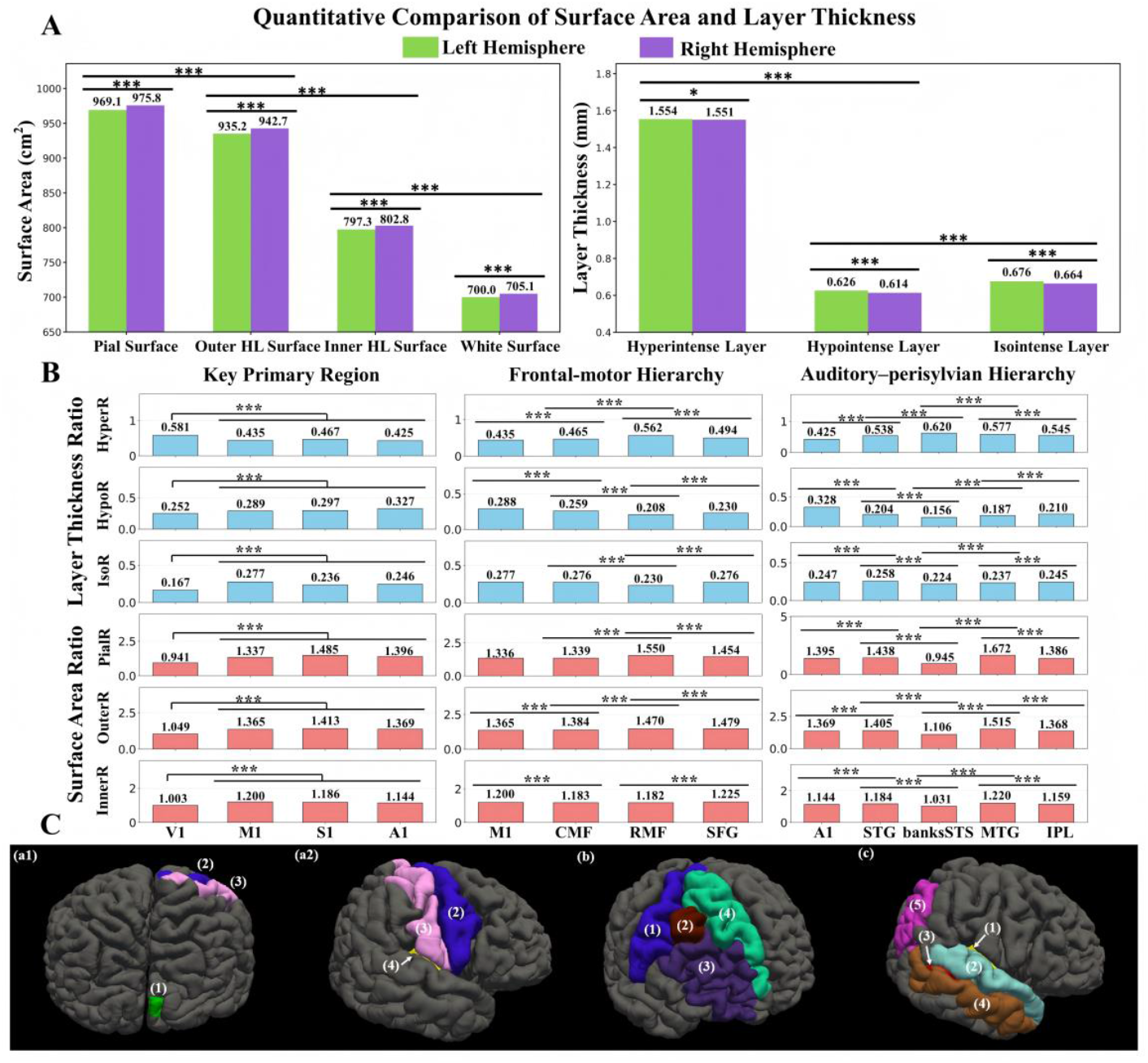
Whole-cortex and regional signal-defined intracortical layer morphometrics across the prespecified cortical organizational framework. (A) Whole-cortex analysis of intracortical layer thickness and surface area; HL denotes the hypointense layer. Inter-hemispheric differences are assessed using paired two-tailed *t*-tests (uncorrected). Within-hemisphere layer or surface comparisons use pairwise paired two-tailed *t*-tests (uncorrected). (B) Regional analysis of thickness ratios (HyperR, HypoR, IsoR) and surface area ratios (PialR, OuterR, InnerR) within: (i) key primary regions, (ii) frontal-motor hierarchy, and (iii) auditory-perisylvian hierarchy. Effects are tested with linear mixed-effects models, followed by Bonferroni-corrected post hoc pairwise comparisons within each system. (C) The cortical organizational framework projected onto 3D pial surface renderings: (a1), (a2) primary regions (V1, M1, S1, A1); (b) frontal-motor hierarchy (M1, CMF, RMF, SFG); (c) auditory-perisylvian hierarchy (A1, STG, banksSTS, MTG, IPL). Asterisks denote significance: *: *P* < .05, **: *P* < .01, ***: *P* < .001.

Layer thickness differed across layers and between hemispheres. The hypointense layer (left: 0.629±0.101 mm; right: 0.614±0.101 mm) was thinner than both the hyperintense (left: 1.554±0.106 mm; right: 1.551±0.111 mm) and isointense layers (left: 0.676±0.074 mm; right: 0.664±0.073 mm) in each hemisphere (all *P* < .001). Across all three layers, thickness was greater in the left hemisphere than in the right (*P*= .02 for the hyperintense layer; *P*< .001 for the hypointense and isointense layers).

In the surface area analysis, all four surfaces were larger in the right hemisphere than in the left (all *P* < .001). Within each hemisphere, surface area decreased monotonically from the pial surface to the white-matter surface, with all pairwise differences between adjacent surfaces reaching *P* <.001.

### 4. Regional Differences Within Three Cortical Systems

In the primary dataset, linear mixed-effects models were used to assess the effects of hemisphere and cortical region on layer-specific thickness ratios (HyperR, HypoR, IsoR) and surface area ratios (PialR, OuterR, InnerR) within three cortical systems: key primary regions, a frontal-motor hierarchy, and an auditory-perisylvian hierarchy (Figure 5B, Table 4). The spatial distribution of the brain regions comprising the three cortical systems is visualized on whole-brain 3D surface renderings in Figure 5C. The selected brain regions and their corresponding Desikan-Killiany (DK) labels are listed in Appendix Table 7.

**Table 4:**
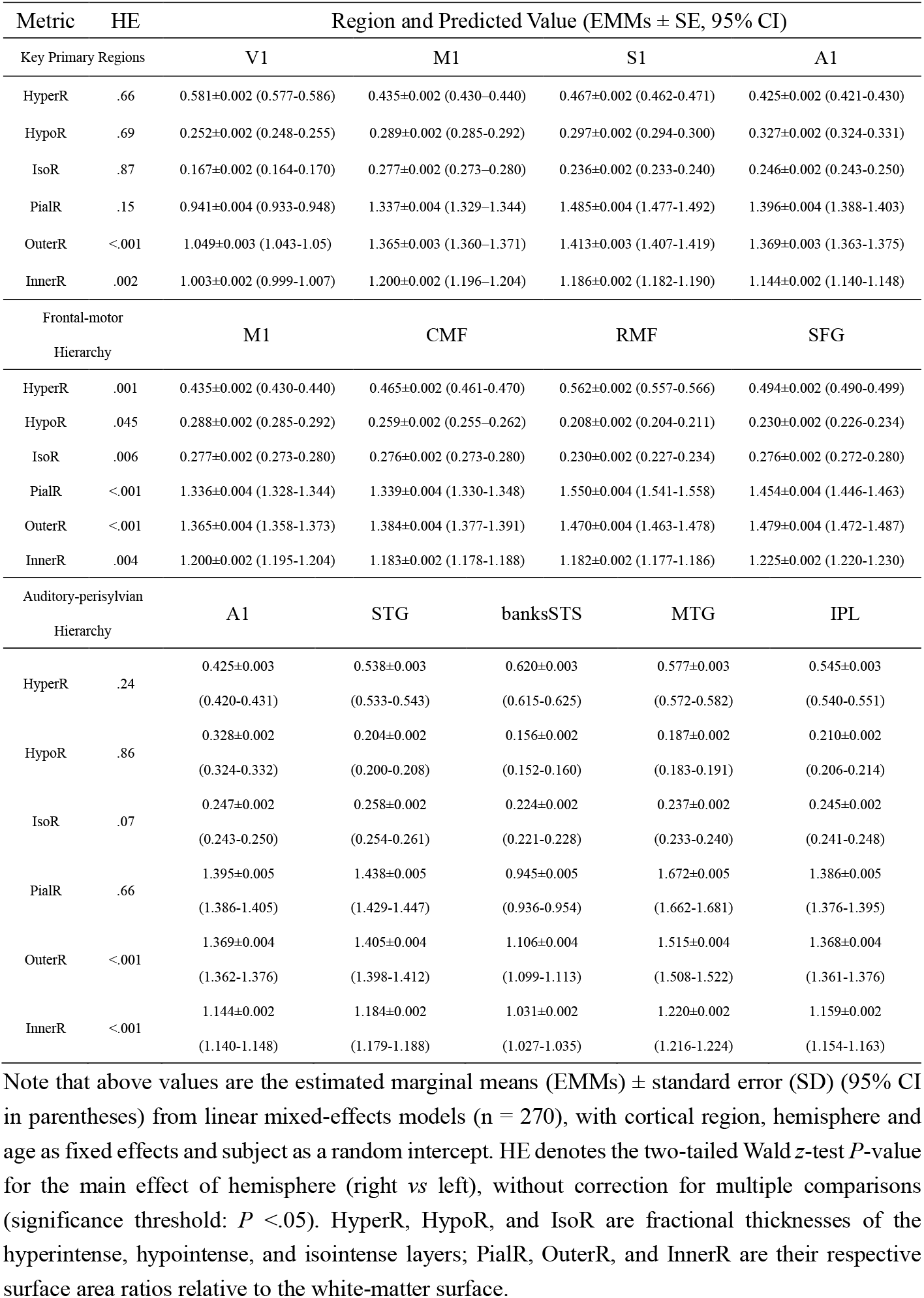
Estimated Marginal Means of Layer-Specific Morphometric Ratios Across Cortical Regions Within Each Cortical System, with Hemisphere Effects (HE)

Key primary regions included the primary visual cortex (V1), primary motor cortex (M1), primary somatosensory cortex (S1), and primary auditory cortex (A1) (Figure 5C (a1) and (a2)). Hemispheric differences were observed for two surface area ratios: OuterR (*P* < .001) and InnerR (*P* .002). For all other metrics, we found no evidence of hemispheric differences (all *P*> .05). Across regions, V1 had higher HyperR than M1, S1, or A1 (all *P* < .001), and lower HypoR and IsoR (all *P* < .001). V1 also exhibited the lowest PialR, OuterR, and InnerR among the four regions (all *P* < .001). (Appendix Table 2)

A frontal-motor hierarchy comprised M1, caudal middle frontal gyrus (CMF), rostral middle frontal gyrus (RMF), and superior frontal gyrus (SFG) (Figure 5C (b)). All metrics showed hemispheric differences (all *P* <.05). Regionally, HyperR increased from M1 to RMF and decreased slightly in SFG (all consecutive pairwise *P* < .001). In contrast, HypoR and IsoR decreased from M1 to RMF and then increased toward SFG; all consecutive pairwise comparisons reached *P* < .001, except for IsoR between M1 and CMF (*P* >.99). For surface area ratios, OuterR increased stepwise along the hierarchy (all consecutive pairwise *P* < .001). PialR rose from M1 to RMF and declined slightly in SFG, whereas InnerR decreased from M1 to RMF and increased in SFG. All consecutive pairwise differences reached *P* < .001, except for M1 versus CMF in PialR (*P* >.99) and CMF versus RMF in InnerR (*P* >.99). (Appendix Table 3)

An auditory-perisylvian hierarchy included A1, superior temporal gyrus (STG), banks of the superior temporal sulcus (banksSTS), middle temporal gyrus (MTG), and inferior parietal lobule (IPL) (Figure 5C (c)). Hemispheric effects were observed only for OuterR and InnerR (both *P*< .001); for all other metrics, we found no evidence of hemispheric asymmetry (all *P* >.05). Regionally, HyperR increased from A1 to STG, peaked in banksSTS, and declined through MTG to IPL, whereas HypoR decreased to a minimum in banksSTS and then increased toward IPL (all consecutive pairwise *P* < .001). A similar pattern was observed for IsoR, PialR, OuterR, and InnerR: values rose from A1 to STG, dropped to a minimum in banksSTS, increased in MTG, and declined in IPL (all consecutive pairwise *P* < .001). (Appendix Table 4)

### 5. Potential Clinical Utility of BrainMLSR-Derived Metrics

We conducted a proof-of-concept analysis of intracortical metrics in Heschl’s gyrus in a 1:1 matched cohort of 19 patients with left TLE and 19 healthy controls (HC) from the primary dataset, pairwise matched for age (exact or nearest year) and, where possible, sex.

Total cortical thickness was lower in TLE than HC in both hemispheres (left: 2.135±0.740 *vs* 2.670±0.192 mm; right: 2.268±0.931 *vs* 2.906±0.221 mm; both *P*<.001), driven by reduced hypointense and isointense layer thickness, whereas hyperintense layer thickness was relatively preserved (Appendix Table 6). Consistent with this pattern, HyperR was higher in TLE versus HC (left: 0.523±0.085 *vs* 0.417±0.036; right: 0.520±0.083 *vs* 0.412±0.050; both *P* < .001), with corresponding reductions in HypoR and IsoR (Appendix Table 5). Additional surface-based ratios are reported in Appendix Table 5. Representative examples (Figure 6A) illustrate higher HyperR alongside reduced total cortical thickness in TLE.

**Figure 6.**
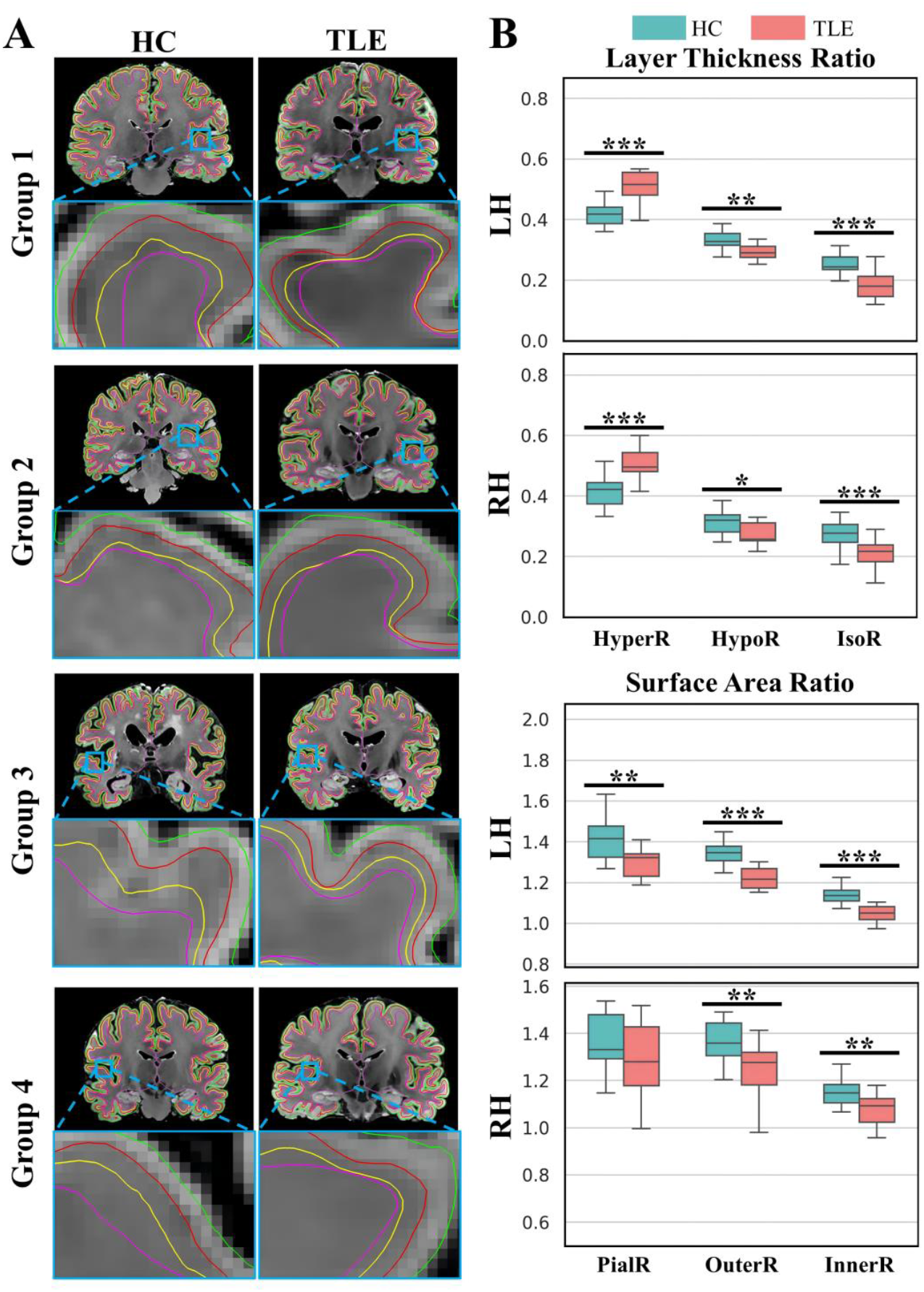
Signal-defined intracortical layer morphometrics in Heschl’s gyrus of left temporal lobe epilepsy (TLE). (A) Representative 3D FLAIR images from healthy controls (HC) and temporal lobe epilepsy (TLE) patients are shown for four groups: groups 1 and 2 display left-hemisphere examples, while groups 3 and 4 show right-hemisphere examples. (B) Quantitative comparison of TLE versus HC in three cortical thickness ratios (HyperR, HypoR, IsoR) and three surface area ratios (PialR, OuterR, InnerR). Group differences are assessed using two-tailed Mann-Whitney U tests without multiple comparison correction. Significant differences are indicated by horizontal lines: *: *P* < .05, **: *P* < .01, ***: *P* < .001.

## DISCUSSION

Standardized *in vivo* whole-cortex quantification of signal-defined intracortical layering from routinely acquired structural MRI remains limited, constraining translational applications of intracortical layer morphometrics. In this study, using a single 5T 3D T2-weighted FLAIR acquisition, we developed BrainMLSR, an automated multi-layer surface reconstruction framework, to delineate three signal-defined intracortical layers across the cerebral cortex *in vivo* (n=270). High agreement with manual annotations (Dice similarity coefficient, 0.960 ± 0.003), submillimeter test-retest reliability (average symmetric surface distance, <0.1 mm; Hausdorff distance, <1.4 mm), and cross-site feasibility (Dice similarity coefficient, 0.954 ± 0.009) supporting BrainMLSR-derived measures as repeatable participant-level readouts.

Prior approaches define an intracortical coordinate system with equidistant or equivolume models; these are interpolated depth surfaces, not the signal-defined boundaries (11,22). As a result, depth-model approaches cannot directly quantify *in vivo* whether an intracortical band shifts radially, changes in boundary sharpness, or shows altered between-layer contrast, which may occur even when total cortical thickness is preserved. Multi-inversion recovery MRI can generate T1-sensitive intracortical profiles that are clustered to delineate lamination patterns but requires multiple acquisitions and longer protocols (7,23), and T1w/T2w myelin maps contain B1+ transmit-field biases that can yield spurious associations without correction (24). In contrast, BrainMLSR reconstructs signal-defined surfaces from a single routinely interpretable structural contrast, and enables participant-level, signal-based morphometry without additional multi-contrast protocols.

We used HyperR, HypoR, and IsoR to summarize the relative radial partitioning across the pial-to-gray-white span *in vivo*. Across the prespecified cortical organizational framework, these thickness proportions showed consistent region- and system-dependent differences, indicating that signal-defined intracortical partitioning is not radially uniform across the cortex *in vivo*. To move beyond isolated areal contrasts, we examined two prespecified unimodal-to-association or transmodal hierarchies (fronto-motor and auditory-perisylvian) and observed ordered profile shifts with localized deviations along the prespecified ROI ordering (25,26).

In parallel, PialR, OuterR, and InnerR provided tangential information complementary to layer thickness and were sensitive to local folding geometry. Consistent with evidence that cortical thickness and surface area are shaped by partly dissociable genetic and developmental influences, these radial and tangential measures did not necessarily vary in parallel across regions (27).

Within the fronto-motor hierarchy, PialR aligned with thickness patterns at key nodes, whereas OuterR and InnerR followed specific trends (i.e., a monotonic anterior increase in OuterR), indicating node- and boundary-dependent concordance versus divergence between radial redistribution and tangential scaling. In the auditory-perisylvian hierarchy, banksSTS formed aturning point where HyperR and HypoR showed a reciprocal peak-trough pattern (HyperR peak, HypoR nadir), coinciding with lower surface area ratios across surfaces. Because banksSTS lies on the posterior superior temporal sulcus (STS) at the interface between the superior and middle temporal cortices, and the posterior STS is implicated in audiovisual or multisensory associative processing (28-30), this inflection may suggest that signal-defined intracortical layer morphometric metrics are sensitive to a functionally relevant anatomic transition zone along the perisylvian axis. Meanwhile, sulcal-bank geometry and atlas-defined parcel borders may also contribute to the observed feature. Within the primary system, V1 differed from other primary cortices in all metrics, supporting that substantial signal-defined radial differences can exist among unimodal areas. One possible explanation is that the granular and highly myelinated characteristics of V1 provide a neuroanatomically plausible context for such prominent banding-related differences (10,31,32). Collectively, these system-scale patterns support BrainMLSR as a standardized *in vivo* readout of signal-defined intracortical layer morphometrics on 5T 3D FLAIR and provide a normative reference for translational applications.

To demonstrate potential clinical utility, we performed a proof-of-concept analysis in left TLE without additional MRI-visible parenchymal lesions on review (except hippocampal sclerosis). Bilateral Heschl’s gyrus showed shifts in signal-defined radial partitioning, driven by disproportionate thinning of hypointense and isointense layers with preservation of the hyperintense layer, which may complement FLAIR assessment and warrants validation in larger cohorts.

This study has several limitations. First, intracortical FLAIR contrast reflects a composite of tissue water, myelin-related relaxation, and perivascular effects; thus, reconstructed layers represent *in vivo* imaging-derived measures rather than direct histologic laminae or a single microstructural mechanism. *Ex vivo* validation, quantitative mapping, and multimodal imaging would improve biological interpretability. Second, despite support from manual annotation, test-retest reliability, age-adjusted analysis, and cross-site feasibility, the main cohort was single-center, limiting generalizability across centers, scanners, protocols, and populations. Third, the three-layer model reflects the most stable partition achievable with current 5T FLAIR; finer segmentation will require higher resolution, improved signal-to-noise ratio, and advanced reconstruction algorithms.

In summary, Brain Multi-Layer Surface Reconstruction (BrainMLSR) automatically provides accurate and repeatable *in vivo* reconstruction of three signal-defined intracortical layers from a single 5T 3D T2-weighted FLAIR acquisition, enabling whole-cortex, region-dependent boundary-based morphometry whose coherent system-level organization along prespecified primary regions and unimodal-to-association (transmodal) hierarchies supports an interpretable neuroanatomical context for future translational and longitudinal studies.

## Abbreviations

HyperR: Hyperintense Layer Thickness Ratio
HypoR: Hypointense Layer Thickness Ratio
IsoR: Isointense Layer Thickness Ratio
PialR: Pial Surface Area Ratio
OuterR: Outer Hypointense Layer Surface Area Ratio
InnerR: Inner Hypointense Layer Surface Area Ratio

