## supplemental material for "*In Vivo* Surface Reconstruction of Three Signal-defined Intracortical Layers Using 5T 3D FLAIR MR Imaging"

### Appendix

#### Appendix S1

This study included three healthy participant datasets, all evaluated using 5T MRI (uMR Jupiter, United Imaging Healthcare) and subject to the same eligibility workflow (Figure 1A)). Participants were initially identified from 5T brain MRI scans acquired during site-specific accrual periods. All participants (age between 18 and 80 years) were prescreened using 3D T2-weighted FLAIR images.

Final eligibility required the absence of any of the following exclusion criteria: (1) duplicate scans from the same individual (only the first or highest-quality session retained); (2) key sequence noncompliance, defined as isotropic voxel size >1.0 mm, incomplete cerebral cortical coverage, or deviation from prescribed TR/TE/TI parameters; (3) any congenital or acquired intracranial structural abnormality; (4) history of stroke, brain tumor, traumatic brain isnjury, or prior intracranial surgery (regardless of indication); (5) documented or self-reported diagnosis of epilepsy, Parkinson disease, or multiple sclerosis; (6) severe global brain atrophy on visual inspection or quality control failure due to severe motion artifacts, insufficient anatomical coverage, or surface-laminar reconstruction failure (e.g., non-convergence or topological defects).

Primary dataset: A consecutive series of imaging sessions was identified from Qilu Hospital (Qingdao) of Shandong University (Center 1), between February and July 2024. From 300 initially screened sessions, 270 met all eligibility and quality criteria and were included in the primary analysis (146 men, 124 women; mean age: 54.4±14.5 years; range: 18-78 years). Four participants (a 52-year-old man, 23-year-old woman, a 60-year-old woman, and a 45-year-old man) were selected for expert manual annotation to assess reconstruction fidelity of our method.

Cross-site dataset: To evaluate the feasibility of the model across imaging sites, two participants (a 68-year-old man and a 61-year-old woman) were recruited from Shandong Provincial Third Hospital (Center 2) in August 2025. Center 2 also utilized a 5T MRI (uMR Jupiter, United Imaging Healthcare), and the participants met all core eligibility criteria outlined above. The two participants were selected for expert manual annotation to assess reconstruction fidelity under cross-site conditions.

Test-retest dataset: To assess test-retest reliability, 13 additional participants were prospectively enrolled at Center 1 between March and July 2025. Each underwent two identical 5T MRI scans separated by a ~15-minute interval using the same scanner and imaging protocol. All participants satisfied the core eligibility criteria above (6 men, 7 women; mean age: 48.2±14.3 years; range: 25-67 years).

Temporal lobe epilepsy dataset: To evaluate potential clinical applicability, we conducted a proof-of-concept retrospective analysis of an ongoing prospectively enrolled cohort of consecutive hospitalized patients with unilateral temporal lobe epilepsy (TLE) in the Neurology Department of Qilu Hospital (Center 1). Enrollment began in March 2024 and is ongoing. The clinical diagnosis of unilateral TLE followed International League Against Epilepsy criteria, and was determined by epileptologists using a comprehensive evaluation that included video-EEG telemetry, seizure semiology, neuropsychological assessment, and neuroimaging. For each patient, neuroimaging was reviewed jointly by at least one radiologist and one neurologist. Seizure onset was lateralized to the left temporal lobe on the basis of this comprehensive evaluation, and only patients with left temporal lobe epilepsy were included in the present analysis. As part of the prospective study protocol, all patients underwent 5T brain MRI (uMR Jupiter; United Imaging Healthcare), including 3D T1-

weighted imaging and T2-weighted FLAIR, MR spectroscopy, with or without glutamate chemical exchange saturation transfer imaging. For the present BrainMLSR analysis, we retrospectively selected participants with left temporal lobe epilepsy who were scanned between January and June 2025 with interictal 5T 3D FLAIR available (no clinically apparent seizures during the scan) and sufficient image quality for surface reconstruction. Additional inclusion criteria required absence of other brain or psychiatric diseases and right-handedness. Exclusion criteria were brain parenchymal lesions on MRI review other than hippocampal sclerosis (e.g., tumor or vascular malformation) and poor image quality (including substantial motion artifact or incomplete cortical coverage) (Figure 1B). The seizure-onset side was defined by video-EEG telemetry, and the contralateral side was defined as the opposite hemisphere.

This selection yielded 19 patients with left TLE (10 women; mean age, 40.9±15.5 years). 19 healthy controls were selected from the original healthy dataset to match the TLE group by age (exact or nearest year) and, when possible, sex; all 19 pairs were age matched and 17 pairs were sex matched, resulting in 19 controls (11 men, 8 women). This retrospective analysis was conducted under institutional review board approval (Qilu Hospital, No. KYLL-KS-2025005). Written informed consent was obtained from all participants.

No participant overlapped with any previously published or in-press study from our group. The sample size of the primary dataset was determined by the number of eligible, high-quality scans available during the accrual period, and is consistent with dataset sizes reported in contemporary high-field MRI studies.

#### Appendix S2

All MRI examinations at Center 1 were performed using a 5T MRI scanner (uMR Jupiter; United Imaging Healthcare). No sedatives or contrast agents were administered before scanning. A high-resolution 3D T2-weighted FLAIR sequence was used with the following parameters: repetition time (TR), 6500ms; echo time (TE), 370ms; inversion time (TI), 1950ms; flip angle, 61°; matrix size, 412 × 330 × 480; and isotropic voxel size, 0.533 × 0.533 × 0.533mm$^3$. Consistent scanning parameters were applied across all participants, including those in the test-retest dataset.

At Center 2, 3D T2-weighted FLAIR images were included for the cross-site dataset using the following parameters: TR, 6000ms; TE, 335.34ms; TI, 1855ms; flip angle, 78°; matrix size, 550 × 550 × 320; and voxel size, 0.4 × 0.4 × 0.8mm$^3$.

#### Appendix S3

The initialization of the inner and outer boundaries of the hypointense cortical layer is performed using signal intensity profiles derived from 5T 3D T2-weighted FLAIR MR images. This procedure relies on an initial pair of topologically consistent triangular mesh surfaces representing the white matter (inner) and pial (outer) cortical boundaries, for which corresponding vertices are in one-to-one spatial correspondence. For each corresponding vertex pair on the inner and outer cortical surfaces, a straight sampling path is constructed through the cortical ribbon. Along this path, 100 equidistant points are sampled from the inner (white matter) surface to the outer (pial) surface, and voxel intensities at these locations are interpolated from the MRI volume to generate a one-dimensional intensity profile spanning the full cortical depth, shown as the solid line in Figure 2(c). The corresponding gradient profile is then computed by taking the first-order derivative of this intensity profile, illustrated as the dashed line in Figure 2(c).

In 5T 3D T2-weighted FLAIR images, the cortex consistently exhibits a three-layer signal pattern: proceeding from the white matter toward the pial surface, the signal transitions through an isointense layer, followed by a hypointense layer, and finally a hyperintense layer, as illustrated in Figure 2(b). Correspondingly, the sampled intensity profiles typically display a characteristic morphology: intensity first decreases to a minimum within the hypointense layer and then rises to the hyperintense layer, consistent with the representative intensity profile shown in Figure 2(c).

Based on this reproducible signal trend, the boundaries of the hypointense layer are initialized using gradient-based criteria. The outer surface of the hypointense layer, defined as the interface between the hypointense and hyperintense layers, corresponds to the onset of the rapid signal increase observed in the intensity profile along the sampling path. This point is identified on the gradient profile as the location where the intensity gradient is positive and attains its maximum magnitude, which corresponds to the steepest ascent into the hyperintense layer on the intensity profile (Figure 2(c)). The inner surface of the hypointense layer, defined as the interface between the isointense and hypointense layers, is constrained to lie between the white-matter surface and the previously-identified outer surface of the hypointense layer. Within this interval, it corresponds to the onset of the most rapid signal decline in the intensity profile along the sampling path. This point is identified on the gradient profile as the location where the intensity gradient is negative and attains its minimum magnitude, which corresponds to the steepest descent into the hypointense layer on the intensity profile (Figure 2(c)).

These gradient-derived estimates provide anatomically-plausible initial surfaces for the hypointense layer, which are subsequently jointly refined through the energy function-based optimization framework of BrainMLSR.

#### Appendix S4

To jointly optimize the coarse initial surfaces, we introduce a multi-surface optimization strategy with the energy function defined below. By iteratively minimizing this energy function, the inner and outer surfaces of the hypointense layer are deformed and refined, ultimately producing anatomically plausible and well-aligned hypointense layer surfaces.

$$\begin{aligned} min\, E(v_{1,i}, v_{2,i}) = &\sum_{k=1,2} \alpha_k \sum_i S_{\text{surface}}(v_{k,i}) \\ &+ \sum_{k=0,1,2} \beta_k \sum_i (D_{\text{vertex}}(v_{k,i}) - D_{\text{adjacent}}(v_{k,i}))^2 \\ &- \sum_{k=1,2} \gamma_k \sum_i |\nabla I(v_{k,i})|^2, \end{aligned} \quad (1)$$

where $v_{k,i}$ is the $i$-th vertex of the $k$-th surface with $k = 0, 1, 2, 3$ indexing the four surfaces from white matter to pia surface (Figure 2(D1)). $\alpha_k$, $\beta_k$, and $\gamma_k$ are tunable hyperparameters, respectively. $\nabla I$ refers to the gradient of the FLAIR image.

Specifically, to enhance the smoothness of each surface, we introduce a surface smooth constraint as shown in Figure 2(D2), defined as $S_{\text{surface}}(v_{k,i}) = ||v_{k,i} - \frac{1}{|\mathcal{N}(v_{k,i})|}\sum_{v_{k,j}\in\mathcal{N}(v_{k,i})} v_{k,j}||^2$, where $\mathcal{N}(v_{k,i})$ denotes the set of vertices $v_{k,j}$ adjacent to $v_{k,i}$ on the $k$-th surface (Figure 2(D2)).

To enforce local uniform spacing between adjacent surfaces, we introduce a surface distance consistency penalty (Figure 2(D3)). This term minimizes the discrepancy between the direct distance from a vertex $v_{k,i}$ on the current surface to the corresponding vertex $v_{k+1,i}$ on the adjacent surface, formulated as $D_{\text{vertex}}(v_{k,i}) = ||v_{k,i} - v_{k+1,i}||$, and the average distance from the adjacent vertices of the vertex $v_{k,i}$ to the adjacent surface, formulated as $D_{\text{adjacent}}(v_{k,i}) = \frac{1}{|\mathcal{N}(v_{k,i})|}\sum_{v_{k,j}\in\mathcal{N}(v_{k,i})} ||v_{k,j} - v_{k+1,j}||$.

Next, we introduce a fidelity term to ensure that the reconstructed surfaces accurately follow cortical signal-layer boundaries by driving vertices toward regions of high image-gradient magnitude. Finally, we propose a surface intersection constraint that monitors each vertex displacement to prevent geometric intersections between adjacent surfaces (Figure 2(D4)) (as detailed in Appendix S5).

Finally, the following hyperparameters were used in the optimization: $\alpha_{1,2} = 3$, $\beta_{1,2,3} = 1$, $\gamma_{1,2} = 0.5$. The algorithm was run for 80 iterations.

#### Appendix S5

In cortical structure modeling, the surfaces of interest, ordered from innermost to outermost, are the cortical inner (white matter) surface, the inner surface of the hypointense layer, the outer surface of the hypointense layer, and the cortical outer (pial) surface. Due to strict one-to-one vertex correspondence across these four layers, their spatial configuration must adhere to a fixed radial ordering: along any sampling path connecting corresponding vertices from the white matter to pial surfaces, the sequence must be $v_{0,\mathrm{i}}$, $v_{1,\mathrm{i}}$, $v_{2,\mathrm{i}}$, $v_{3,\mathrm{i}}$. Maintaining this anatomical hierarchy during deformation optimization is essential; violations, such as the hypointense layer's inner surface vertex moving beyond its outer surface, lead to surface intersections that contradict the known laminar organization of the cortex (Figure 2 (d)(4)).

To enforce this constraint, we leverage the fact that all deformations of the hypointense layer surfaces are restricted to the straight line between the fixed white matter and pial surface vertices. After each deformation step, we perform a quantitative check of the radial ordering as follows. Let $v_{0,\mathrm{i}}$ and $v_{3,\mathrm{i}}$ denote the fixed positions of the corresponding white matter and pial vertices, and let $v_{1,\mathrm{i}}$ and $v_{2,\mathrm{i}}$ be the current (possibly deformed) positions of the hypointense layer's inner and outer surface vertices. We define three vectors originating from $v_{0,\mathrm{i}}$:

$$v_{\mathrm{pial}} = v_{3,\mathrm{i}} - v_{0,\mathrm{i}},\ v_{\mathrm{outer}} = v_{2,\mathrm{i}} - v_{0,\mathrm{i}},\ v_{\mathrm{inner}} = v_{1,\mathrm{i}} - v_{0,\mathrm{i}} \tag{1}$$

We need to verify that their scalar projections onto the radial direction satisfy:

$$0 < v_{\mathrm{inner}} \cdot v_{\mathrm{pial}} < v_{\mathrm{outer}} \cdot v_{\mathrm{pial}} < v_{\mathrm{pial}} \cdot v_{\mathrm{pial}} \tag{2}$$

This ensures that the vertices of both the inner and outer surfaces of the hypointense layer lie between the cortical inner and outer surface vertices, and that the hypointense layer's inner surface vertex is closer to the cortical inner surface than its outer surface vertex, thereby preserving the correct relative positional ordering of the vertices across all four surfaces. If this condition is not satisfied, it indicates an intersection between surfaces, and the corresponding deformation of the vertex is therefore rejected.

#### Appendix S6

Whole-hemisphere averages of layer-specific cortical thickness (hyperintense layer, hypointense layer and isointense layer) and total cortical thickness were computed separately for the left and right hemispheres. Similarly, surface areas were derived for four reconstructed surfaces: pial surface, outer and inner hypointense layer surfaces, and white-matter surface, each also calculated independently for the left and right hemispheres.

To assess lateralization, paired two-tailed *t*-tests were used to compare each metric between the left and right hemispheres across participants (e.g., left *vs* right superficial thickness). These hemisphere-level comparisons were prespecified and intended as descriptive summaries; therefore, *P* values were interpreted without adjustment for multiple comparisons across metrics. Within each hemisphere, differences among the intracortical layers (for thickness) or among the surfaces (for area) were evaluated using paired two-tailed *t*-tests and no correction for multiple comparisons was applied.

To facilitate comparisons across cortical regions that differ in absolute size, we computed layer-specific morphometric ratios: (1) thickness ratios, defined as the proportion of each cortical layer's thickness relative to total cortical thickness, and (2) surface area ratios, defined as the ratio of the pial, inner hypointense, and outer hypointense surface areas to the white-matter surface area.

For each of the three a priori-defined cortical systems—(i) primary regions (V1, M1, S1, A1), (ii) frontal-motor hierarchy (M1, CMF, RMF, SFG), and (iii) auditory-perisylvian hierarchy (A1, STG, banksSTS, MTG, IPL)—separate linear mixed-effects models were fitted for each of the six layer-specific metrics (three thickness ratios and three surface area ratios). The general model structure was:

$$Ratio \sim Region + Hemisphere + Age + (1 \mid Subject) \tag{2}$$

where Region (categorical) and Hemisphere (left vs right) were included as fixed effects to test regional differences and potential lateralization, respectively. Age was included as a continuous fixed-effect covariate to statistically adjust for its potential confounding influence on morphometric ratios. A random intercept for subject accounted for within-participant correlation across multiple regions.

Estimated marginal means (EMMs) for each region were computed at the sample mean age (54.4 years) and are reported as mean ± standard error (SE) with 95% confidence intervals (CIs). Post hoc pairwise comparisons among regions within each cortical systems were conducted by testing differences between EMMs using two-tailed Wald *z*-tests derived from the linear mixed-effects model's parameter covariance matrix. To control the family-wise error rate, Bonferroni correction was applied separately within each hierarchy based on the number of possible pairwise contrasts.

The effect of hemisphere was assessed solely via the main effect coefficient for Hemisphere in each model (coded as right *vs* left). Given that hemisphere is a binary factor, only a single contrast was tested; therefore, no correction for multiple comparisons was applied. Significance was determined directly from the model's two-tailed Wald *z*-test for the hemisphere term.

To assess clinical relevance, we compared intracortical thickness, surface area, and their corresponding ratios in bilateral Heschl's gyrus, a core auditory cortical region within the temporal lobe, between 19 patients with left temporal lobe epilepsy (TLE) and 19 age- and sex-matched healthy controls (HC). Both hemispheres were analyzed to assess whether group differences were lateralized relative to the seizure-onset side. The Heschl's gyrus ROI corresponds to the “transversetemporal” label in the Desikan-Killiany cortical atlas (see Appendix Table 7). This region was prespecified a priori because it is an anatomically well-defined and consistently labeled temporal lobe parcel that maps to the transverse temporal gyrus (Heschl’s gyrus), which contains primary auditory cortex and is clinically relevant to auditory symptoms that may occur in temporal lobe seizures. Group differences were evaluated using a two-tailed Mann-Whitney U test without correction for multiple comparisons, as the analysis was hypothesis-driven and restricted to this single *a priori* region of interest.

All statistical analyses were conducted using Python (version 3.8; Python Software Foundation) by S.C., and statistical significance for all tests was defined as $P < .05$.

**Appendix Table 1:** Within-Hemisphere Pairwise Comparisons of Signal-Defined Intracortical layer thickness and Surface Area of Reconstructed Cortical Surfaces

| Thickness (LH) | Hyperintense | Hypointense Layer | Isointense Layer | Total Cortex |
|---|---|---|---|---|
| Hyperintense Layer | - | <.001 | <.001 | <.001 |
| Hypointense Layer | - | - | <.001 | <.001 |
| Isointense Layer | - | - | - | <.001 |
| Total Cortex | - | - | - | - |
| **Thickness (RH)** | **Hyperintense** | **Hypointense Layer** | **Isointense Layer** | **Total Cortex** |
| Hyperintense Layer | - | <.001 | <.001 | <.001 |
| Hypointense Layer | - | - | <.001 | <.001 |
| Isointense Layer | - | - | - | <.001 |
| Total Cortex | - | - | - | - |
| **Surface Area (LH)** | **Pial Surface** | **Outer HL Surface** | **Inner HL Surface** | **White Surface** |
| Pial Surface | - | <.001 | <.001 | <.001 |
| Outer HL Surface | - | - | <.001 | <.001 |
| Inner HL Surface | - | - | - | <.001 |
| White Surface | - | - | - | - |
| **Surface Area (RH)** | **Pial Surface** | **Outer HL Surface** | **Inner HL Surface** | **White Surface** |
| Pial Surface | - | <.001 | <.001 | <.001 |
| Outer HL Surface | - | - | <.001 | <.001 |
| Inner HL Surface | - | - | - | <.001 |
| White Surface | - | - | - | - |

Note that $P$ values above are from paired two-tailed $t$-tests comparing all pairs of layers (for thickness) or surfaces (for area) within each hemisphere, without correction for multiple comparison (significance threshold: $P < .05$). Abbreviations: HL = hypointense layer; LH = left hemisphere; RH = right hemisphere.

**Appendix Table 2**: Bonferroni-Corrected Pairwise Comparisons of Intracortical Layer Morphometric Ratios Among Regions in the Key Primary Regions

| Key Primary Regions | | | | | | | | |
|---|---|---|---|---|---|---|---|---|
| | HyperR | | | | PialR | | | |
| | V1 | M1 | S1 | A1 | V1 | M1 | S1 | A1 |
| V1 | - | <.001 | <.001 | <.001 | - | <.001 | <.001 | <.001 |
| M1 | - | - | <.001 | <.001 | - | - | <.001 | <.001 |
| S1 | - | - | - | <.001 | - | - | - | <.001 |
| A1 | - | - | - | - | - | - | - | - |
| | HypoR | | | | OuterR | | | |
| | V1 | M1 | S1 | A1 | V1 | M1 | S1 | A1 |
| V1 | - | <.001 | <.001 | <.001 | - | <.001 | <.001 | <.001 |
| M1 | - | - | <.001 | <.001 | - | - | <.001 | >.99 |
| S1 | - | - | - | <.001 | - | - | - | <.001 |
| A1 | - | - | - | - | - | - | - | - |
| | IsoR | | | | InnerR | | | |
| | V1 | M1 | S1 | A1 | V1 | M1 | S1 | A1 |
| V1 | - | <.001 | <.001 | <.001 | - | <.001 | <.001 | <.001 |
| M1 | - | - | <.001 | <.001 | - | - | <.001 | <.001 |
| S1 | - | - | - | <.001 | - | - | - | <.001 |
| A1 | - | - | - | - | - | - | - | - |

Note that pairwise comparisons among cortical regions are performed using estimated marginal means based on two-tailed Wald tests. *P* values are adjusted for multiple comparisons using the Bonferroni correction (significance threshold: $P < .05$). HyperR, HypoR, and IsoR represent the thickness of the hyperintense, hypointense, and isointense layers, respectively, relative to total cortical thickness. PialR, OuterR, and InnerR denote the surface area of the pial, outer hypointense, and inner hypointense surfaces, respectively, relative to the white-matter surface area.

**Appendix Table 3**: Bonferroni-Corrected Pairwise Comparisons of Intracortical Layer Morphometric Ratios Among Regions in the Frontal-Motor Hierarchy

| Frontal-motor Hierarchy | | | | | | | | |
|---|---|---|---|---|---|---|---|---|
| | HyperR | | | | PialR | | | |
| | M1 | CMF | RMF | SFG | M1 | CMF | RMF | SFG |
| M1 | - | <.001 | <.001 | <.001 | - | >.99 | <.001 | <.001 |
| CMF | - | - | <.001 | <.001 | - | - | <.001 | <.001 |
| RMF | - | - | - | <.001 | - | - | - | <.001 |
| SFG | - | - | - | - | - | - | - | - |
| | HypoR | | | | OuterR | | | |
| | M1 | CMF | RMF | SFG | M1 | CMF | RMF | SFG |
| M1 | - | <.001 | <.001 | <.001 | - | <.001 | <.001 | <.001 |
| CMF | - | - | <.001 | <.001 | - | - | <.001 | <.001 |
| RMF | - | - | - | <.001 | - | - | - | .001 |
| SFG | - | - | - | - | - | - | - | - |
| | IsoR | | | | InnerR | | | |
| | M1 | CMF | RMF | SFG | M1 | CMF | RMF | SFG |
| M1 | - | >.99 | <.001 | >.99 | - | <.001 | <.001 | <.001 |
| CMF | - | - | <.001 | >.99 | - | - | >.99 | <.001 |
| RMF | - | - | - | <.001 | - | - | - | <.001 |
| SFG | - | - | - | - | - | - | - | - |

Note that pairwise comparisons among cortical regions are performed using estimated marginal means based on two-tailed Wald tests. *P* values are adjusted for multiple comparisons using the Bonferroni correction (significance threshold: *P* <.05). HyperR, HypoR, and IsoR represent the thickness of the hyperintense, hypointense, and isointense layers, respectively, relative to total cortical thickness. PialR, OuterR, and InnerR denote the surface area of the pial, outer hypointense, and inner hypointense surfaces, respectively, relative to the white matter surface area.

**Appendix Table 4**: Bonferroni-Corrected Pairwise Comparisons of Intracortical Layer Morphometric Ratios Among Regions in the Auditory-Perisylvian Hierarchy

| Auditory-perisylvian Hierarchy | | | | | | | | | | |
|---|---|---|---|---|---|---|---|---|---|---|
| | HyperR | | | | | PialR | | | | |
| | A1 | STG | banksSTS | MTG | IPL | A1 | STG | banksSTS | MTG | IPL |
| A1 | - | <.001 | <.001 | <.001 | <.001 | - | <.001 | <.001 | <.001 | .79 |
| STG | - | - | <.001 | <.001 | .005 | - | - | <.001 | <.001 | <.001 |
| banksSTS | - | - | - | <.001 | <.001 | - | - | - | <.001 | <.001 |
| MTG | - | - | - | - | <.001 | - | - | - | - | <.001 |
| IPL | - | - | - | - | - | - | - | - | - | - |
| | HypoR | | | | | OuterR | | | | |
| | A1 | STG | banksSTS | MTG | IPL | A1 | STG | banksSTS | MTG | IPL |
| A1 | - | <.001 | <.001 | <.001 | <.001 | - | <.001 | <.001 | <.001 | >.99 |
| STG | - | - | <.001 | <.001 | <.001 | - | - | <.001 | <.001 | <.001 |
| banksSTS | - | - | - | <.001 | <.001 | - | - | - | <.001 | <.001 |
| MTG | - | - | - | - | <.001 | - | - | - | - | <.001 |
| IPL | - | - | - | - | - | - | - | - | - | - |
| | IsoR | | | | | InnerR | | | | |
| | A1 | STG | banksSTS | MTG | IPL | A1 | STG | banksSTS | MTG | IPL |
| A1 | - | <.001 | <.001 | <.001 | >.99 | - | <.001 | <.001 | <.001 | <.001 |
| STG | - | - | <.001 | <.001 | <.001 | - | - | <.001 | <.001 | <.001 |
| banksSTS | - | - | - | <.001 | <.001 | - | - | - | <.001 | <.001 |
| MTG | - | - | - | - | <.001 | - | - | - | - | <.001 |
| IPL | - | - | - | - | - | - | - | - | - | - |

Note that pairwise comparisons among cortical regions are performed using estimated marginal means based on two-tailed Wald tests. *P* values are adjusted for multiple comparisons using the Bonferroni correction (significance threshold: $P < .05$). HyperR, HypoR, and IsoR represent the thickness of the hyperintense, hypointense, and isointense layers, respectively, relative to total cortical thickness. PialR, OuterR, and InnerR denote the surface area of the pial, outer hypointense, and inner hypointense surfaces, respectively, relative to the white matter surface area.

**Appendix Table 5**: Intracortical Layer Thickness and Surface Area Ratios in Heschl's Gyrus in Left Temporal Lobe Epilepsy and Matched Healthy Controls (HC).

| Metric | Hemi | Ratio Type | HC (Mean±SD) | TLE (Mean±SD) | *P* Value |
|---|---|---|---|---|---|
| Thickness Ratio | LH | HyperR | 0.417±0.036 | 0.523±0.085 | <.001 |
| | | HypoR | 0.330±0.030 | 0.284±0.071 | .005 |
| | | IsoR | 0.253±0.037 | 0.193±0.068 | <.001 |
| | RH | HyperR | 0.412±0.050 | 0.520±0.083 | <.001 |
| | | HypoR | 0.312±0.042 | 0.268±0.073 | .02 |
| | | IsoR | 0.276±0.048 | 0.212±0.044 | <.001 |
| Surface Area Ratio | LH | PialR | 1.410±0.096 | 1.294±0.221 | .005 |
| | | OuterR | 1.351±0.067 | 1.224±0.172 | <.001 |
| | | InnerR | 1.140±0.045 | 1.060±0.076 | <.001 |
| | RH | PialR | 1.362±0.117 | 1.253±0.234 | .13 |
| | | OuterR | 1.363±0.089 | 1.228±0.140 | .002 |
| | | InnerR | 1.150±0.056 | 1.076±0.062 | .002 |

Note that the above values are presented as mean ± standard deviation (SD). Between-group comparisons are performed using the two-tailed Mann-Whitney U test (significance threshold: $P < .05$), without correction for multiple comparison. HyperR, HypoR, and IsoR denote the proportions of hyperintense, hypointense, and isointense layer thicknesses relative to total cortical thickness, respectively. PialR, OuterR, and InnerR represent the proportions of pial, outer hypointense, and inner hypointense surface areas relative to the white-matter surface area, respectively. Abbreviations: LH = left hemisphere; RH = right hemisphere.

**Appendix Table 6**: Intracortical Layer Thickness and Surface Area in Heschl's Gyrus in Left Temporal Lobe Epilepsy and Matched Healthy Controls (HC).

| Metric | Hemi | Type | HC (Mean ± SD) | EP (Mean ± SD) | *P* value |
|---|---|---|---|---|---|
| Layer Thickness | LH | Hyper | 1.111 ± 0.090 | 1.110 ± 0.368 | .02 |
| | | Hypo | 0.881 ± 0.102 | 0.619 ± 0.262 | <.001 |
| | | Iso | 0.678 ± 0.128 | 0.406 ± 0.225 | <.001 |
| | | Total | 2.670 ± 0.192 | 2.135 ± 0.740 | <.001 |
| | RH | Hyper | 1.191 ± 0.123 | 1.184 ± 0.528 | .054 |
| | | Hypo | 0.907 ± 0.141 | 0.607 ± 0.310 | <.001 |
| | | Iso | 0.808 ± 0.176 | 0.478 ± 0.232 | <.001 |
| | | Total | 2.906 ± 0.221 | 2.268 ± 0.931 | <.001 |
| Surface Area | LH | Pial | 536.6 ± 95.6 | 423.2 ± 192.1 | .06 |
| | | Outer | 514.4 ± 93.5 | 397.4 ± 177.4 | .04 |
| | | Inner | 434.1 ± 77.4 | 341.5 ± 149.0 | .044 |
| | | White | 380.4 ± 66.2 | 323.4 ± 140.5 | .48 |
| | RH | Pial | 356.9 ± 63.7 | 390.5 ± 291.6 | .64 |
| | | Outer | 356.6 ± 56.8 | 378.4 ± 288.0 | .62 |
| | | Inner | 300.8 ± 45.1 | 336.0 ± 290.5 | .73 |
| | | White | 261.3 ± 34.3 | 314.7 ± 293.7 | .41 |

Note that the above values are presented as mean ± standard deviation (SD). Between-group comparisons are performed using the two-tailed Mann-Whitney U test (significance threshold: $P < .05$), without correction for multiple comparison. Hyper, Hypo, and Iso denote the hyperintense, hypointense, and isointense layers, respectively. Pial, Outer, Inner and White represent the pial, outer hypointense layer, inner hypointense layer and white-matter surfaces, respectively. Abbreviations: LH = left hemisphere; RH = right hemisphere.

**Appendix Table 7**: *A priori* ROI Definitions and Modality/Functional Characterization (DK Labels)

| ROI systems | ROI | DK label (FreeSurfer aparc) | Core anatomic definition | Modality | Functional characterization |
|---|---|---|---|---|---|
| Key primary regions | V1 (primary visual cortex) | pericalcarine | Cortex surrounding the calcarine sulcus on the medial occipital lobe (canonical locus of primary visual cortex). | Unimodal (primary) | Early-stage visual processing (striate cortex / BA17-level function). |
| Key primary regions | M1 (primary motor cortex) | precentral | Precentral gyrus (anterior to the central sulcus), canonical locus of primary motor cortex. | Unimodal (primary) | Voluntary motor execution (primary motor functions). |
| Key primary regions | S1 (primary somatosensory cortex) | postcentral | Postcentral gyrus (posterior to the central sulcus), contains primary somatosensory cortex. | Unimodal (primary) | Somatosensory input processing and integration (S1). |
| Key primary regions | A1 (primary auditory cortex) | transversetemporal | Heschl's gyrus / transverse temporal gyrus on the supratemporal plane; canonical locus of primary auditory cortex. | Unimodal (primary) | Initial cortical sound processing (primary auditory functions). |
| Frontal-motor hierarchy | CMF | caudalmiddlefrontal | Posterior (caudal) portion of the middle frontal gyrus; part of lateral prefrontal cortex territory. | Association (control-related) | Cognitive control / executive processing; overlaps common "frontoparietal control system" lateral PFC component at the systems level. |
| Frontal-motor hierarchy | RMF | rostralmiddlefrontal | Anterior (rostral) portion of the middle frontal gyrus; lateral prefrontal cortex territory (often used to approximate DLPFC-related regions). | Association (control-related) | Working memory / executive control; commonly operationalized within lateral PFC / DLPFC-related targets in cognitive-control literature. |
| Frontal-motor hierarchy | SFG | superiorfrontal | Superior frontal gyrus (dorsal frontal convexity; includes dorsal prefrontal association cortex territories). | Association (control-related; potential transmodal adjacency) | Dorsal prefrontal functions with established coupling to cognitive-control networks (subregion-dependent). |
| Auditory-perisylvian hierarchy | STG | superiortemporal | Superior temporal gyrus, located between the lateral fissure (Sylvian fissure) and superior temporal sulcus. | Unimodal-association (auditory association) | Auditory/speech sound feature processing and phonological computations (auditory association roles). |
| Auditory-perisylvian hierarchy | banksSTS | bankssts | Banks of the superior temporal sulcus (gyral/sulcal lips forming the STS banks). | Heteromodal association (multisensory interface) | Audiovisual/multisensory integration and higher-order speech-related representations (STS literature). |
| Auditory-perisylvian hierarchy | MTG | middletemporal | Middle temporal gyrus on the lateral temporal surface, ventral to the superior temporal gyrus. | Heteromodal association (semantic/transmodal leaning) | Semantic/lexical comprehension and semantic cognition (temporal association cortex). |
| Auditory-perisylvian hierarchy | IPL | inferiorparietal | Inferior parietal cortex on the lateral parietal convexity, posterior to the postcentral gyrus and adjacent to the intraparietal sulcus region. | Heteromodal association / transmodal-related | Multimodal integration and higher-order functions (language/spatial/learning); a canonical parietal component in frontoparietal control system descriptions. |

Note that regions of interest (ROIs) are anatomically defined using the Desikan-Killiany cortical parcellation as implemented in FreeSurfer (aparc annotation), which labels 34 gyral-based cortical parcels per hemisphere. For readability, selected DK parcels are referred to using conventional neuroanatomical shorthand that reflects their canonical location: V1 for the pericalcarine parcel, M1 for precentral, S1 for postcentral, and A1 for transverse temporal.
